# Phase separation-based visualization of protein-protein interactions and kinase activities in plants

**DOI:** 10.1101/2022.09.06.506782

**Authors:** Alaeddine Safi, Wouter Smagghe, Amanda Gonçalves, Ke Xu, Ana Ibis Fernandez, Benjamin Cappe, Franck B. Riquet, Evelien Mylle, Daniël Van Damme, Danny Geelen, Geert De Jaeger, Tom Beeckman, Jelle Van Leene, Steffen Vanneste

## Abstract

Protein activities depend heavily on protein complex formation and dynamic post-translational modifications, such as phosphorylation. Their dynamic nature is notoriously difficult to monitor in planta at cellular resolution, often requiring extensive optimization and high-end microscopy. Here, we generated and exploited the SYnthetic Multivalency in PLants (SYMPL)-vector set to study protein-protein interactions (PPIs) and kinase activities in planta based on phase separation. This technology enabled easy detection of inducible, binary and ternary protein-protein interactions among cytoplasmic, nuclear and plasma membrane proteins in plant cells via a robust image-based readout. Moreover, we applied the SYMPL toolbox to develop an in vivo reporter for SnRK1 kinase activity, allowing us to visualize tissue-specific, dynamic SnRK1 activation upon energy deprivation in stable transgenic Arabidopsis plants. The applications of the SYMPL cloning toolbox lay the foundation for the exploration of PPIs, phosphorylation and other post-translational modifications with unprecedented ease and sensitivity.

## Main

Protein-protein interactions (PPI) and post-translational modifications (PTMs) such as phosphorylation are two cornerstones in the refined signaling networks and processes that control growth and development in all living organisms. The emergence of sensitive mass spectrometry-based proteomics methods has boosted the rapid identification of protein interactors and post-translational modifications (PTMs) for a protein of interest (POI), often yielding large lists of candidates that require further validation ^1^. Validation of identified PPIs and PTMs remains a major technical challenge with traditional, time-consuming and demanding biochemical and microscopic approaches ^2 3 4^. This illustrates the urgent need for the implementation and development of new analytical methods for the rapid and robust *in planta* detection of PPIs and PTMs ^5^.

An interesting physical phenomenon that occurs upon multivalent interaction between biomolecular components is the formation of dense condensates of the interacting components, commonly referred to as liquid-liquid phase separation ^6^. Phase separation relies on the presence of intrinsically disordered regions in proteins and occurs naturally in biological systems, such as stress granules, nucleoli and Cajal bodies ^7 8 9^. Furthermore, it was shown to control numerous physiological processes such as immunity, flowering, auxin signaling, seed germination, CO2 sensing, thermotolerance, endocytosis, transposon silencing and RNA processing ^10 11 12 13 14 15^. For example, AUXIN RESPONSIVE FACTORs (ARFs) undergo phase separation through their intrinsically disordered PB1 domains leading to the formation of a large complex unable to access the nucleus. This nucleo-cytoplasmic partitioning attenuates auxin responsiveness in tissues where it is less needed ^16^. Employing a different mechanism, the transcription factor TERMINATING FLOWER forms nuclear condensates upon sensing reactive oxygen species to repress *ANANTHA* transcription and thus flowering ^17^. In each of these examples, bright fluorescent droplets form upon phase separation of fluorescent protein fusions, enabling easy detection at a high signal-to-noise ratio by standard confocal microscopy.

Phase transition depends on a threshold of protein concentration, temperature, valency and affinity of the individual interaction partners ^18^. These physical principles were only recently exploited for developing analytical sensors of PPIs (Separation of Phase-based Protein Interaction Reporter, SPPIER) or phosphorylation events (Separation of Phase-based Activity Reporter of Kinase, SPARK) in human cell lines ^19 20 21^. These methods use high-affinity, homo-oligomerizing alpha-helical monomers as tags (HOTags/HTs), which allowed for instance the formation of rapamycin-inducible, synthetic multivalency between FRB and FKBP in human cell lines ^19^. In combination with a fluorescent tag, large phase-separated condensates were detected as bright fluorescent droplets in the cell that can be easily detected using a simple microscopic set-up. Here, we transferred this HOTag-dependent technology to plants through the generation of a SYnthetic Multivalency in PLants (SYMPL) toolbox, allowing the rapid development of robust plant PPI and kinase activity reporters. We demonstrate its utility in plants through the detection of a variety of PPIs, covering both binary or ternary PPIs that occur either in the cytoplasm, the nucleus or at the plasma membrane. Finally, we applied SYMPL to develop a readout for the *in vivo* activity of the SNF1-related kinase 1 (SnRK1), a key metabolic regulator ^22^, revealing the dynamics of SnRK1 activation in a growing root. Together, we showcase how phase-separation based on synthetic multivalency represents a novel, easy-to-use analytical method for the *in planta* detection of PPI and PTMs.

## Results

### Phase separation detects direct binary and ternary protein-protein interactions

We first developed a cloning toolbox (SYMPL) for a rapid and flexible fusion of fluorescent HOTags (HTs) to the proteins of interest, that is compatible with the widely used Gateway and Golden Gate cloning strategies (Fig. 1a and Supplementary Table 1). As a first proof-of-concept of SPPIER for monitoring PPIs in plant cells, we tested the well-established direct binary interactions between 1) the core cell cycle regulator CDKA;1 and its scaffold protein CKS1 ^23 24 25^, and 2) the TPLATE and TML subunits of the hetero-octameric endocytic TPLATE complex (TPC) ^26^. As a negative control for TPLATE, we used its reported indirect interaction with the TPC subunit LOLITA ^27^. The components of each binary combination were fused respectively to either mScarlet-HOTag3 and EGFP-HOTag6 or EGFP-HOTag3 and mScarlet-HOTag6, and were transiently overexpressed in *N. benthamiana* leaves. Co-expression of the corresponding constructs resulted in the formation of bright, dual-labeled intracellular droplets for CKS1-CDKA;1 and TPLATE-TML, clearly differing from the localization pattern of the individually expressed constructs (Fig. 1b-c). In contrast, no yellow droplets were observed for the LOLITA-TPLATE combination (Fig. 1d), which is consistent with their reported indirect interaction. Another TPC subunit, TASH3, physically connects TPLATE to LOLITA ^27 28^. Co-expressing TASH3-TagBFP2 as a bridging factor resulted in the formation of, triple-labeled droplets when the three components were co-expressed in the same cell (Fig. 1d and Supplementary Fig. 1a-e), corroborating the ternary interaction between these three TPC subunits. These data demonstrate that SPPIER potently discriminates between direct and indirect PPIs in plants, and that it can be easily adapted to detect multiprotein PPIs.

**Fig. 1.**
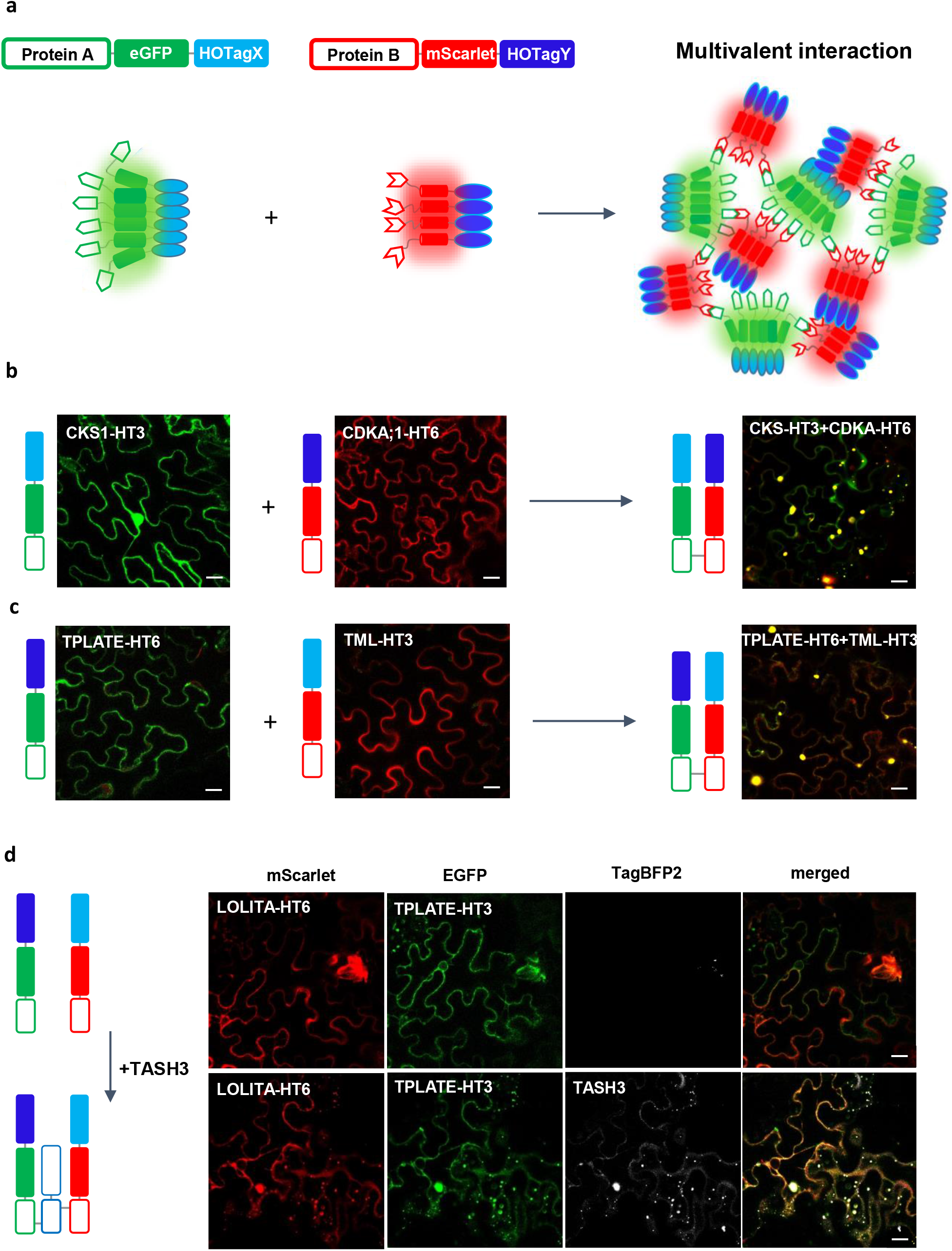
Phase separation to monitor simple and higher-order PPI in plants. **a**, Schematic illustration of phase separation based on the co-expression of two proteins tagged with distinct fluorescently labeled HOTags. Individually expressed HOTagged proteins are homogeneously distributed in the cell. Interaction between protein A and protein B triggers multivalent interaction and liquid-liquid phase-separation of the complexes into bright fluorescent droplets in the cell. **b-c**, Transient co-expression of different HOTag constructs in *N. benthamiana* leaves. Co-expression of CDKA;1-mScarlet-HOTag3 (CDKA;1-HT3) with CKS1-EGFP-HOTag6 (CKS1-HT6) **(b)** or TPLATE-EGFP-HOTag3 (TPLATE-HT3) with TML-mScarlet-HOTag6 (TML-HT6) **(c)** yields the formation of large, dual colored droplets. **d**, Co-expression of LOLITA-mScarlet-HOTag6 (LOLITA-HT6) and TPLATE-HT3 without (upper panel) or with (lower panel) TASH3-TagBFP2. Droplets were only formed in the presence of TASH3-TagBFP2. n=nucleus. Scale bars = 20 μm.

### Simultaneous detection of homo- and hetero-multimerization in the cytoplasm and nucleus

To further expand the palette of proteins that can be assayed by SYMPL, we explored the well-known heteromeric interaction between the GRAS transcription factors SHORTROOT (SHR) and SCARECROW (SCR) upon transient expression in *N. benthamiana* ^29 30^. These transcription factors were shown to act in various complexes that govern root cell identity specification ^31^. Consistently with previous reports ^32 33^, SCR-EGFP localized exclusively to the nucleus (Fig. 2a), whereas SHR-mScarlet was detected both in the nucleus and the cytoplasm (Fig. 2b). Upon co-expression of both fusion proteins, SCR-EGFP sequestered all SHR-mScarlet in the nucleus (Fig. 2c). It remains, however, unknown if the nuclear recruitment of SHR occurs via direct or indirect interaction with SCR. To further explore the underlying molecular mechanism, we co-expressed SHR-mScarlet-HOTag6 and SCR-EGFP-HOTag3 and found that all SHR co-localized with SCR in intensely fluorescent droplets (Fig. 2f), suggesting a direct SHR-SCR interaction. Notably, in contrast to the SCR-EGFP x SHR-mScarlet combination, the intense SHR-mScarlet-HOTag6 x SCR-EGFP-HOTag3 droplets formed in the cytoplasm, suggesting that this is the location where the SHR-SCR interaction is established and that the HOTag-induced phase separation interferes with the subsequent nuclear import of the SHR-SCR complex. Thus, this seems to emphasize a step in the SCR-SHR complex dynamics that was not detectable before. As controls, we analyzed the localization patterns of the individually expressed HOTag fusions. SCR-EGFP-HOTag3 was exclusively nuclear, with the occasional formation of nuclear droplets (Fig. 2d), whereas SHR-mScarlet-HOTag6 appeared mainly in cytoplasmic droplets (Fig. 2e), further corroborating homo-oligomerization of a fraction of these proteins, which was recently observed in the cortex-endodermis initials and the quiescent center cells by high-end microscopy ^34^. These controls illustrate that SHR and SCR can homo-oligomerize in the cytoplasm and nucleus, respectively. Consequently, both control experiments suggest that it is possible to induce phase separation with only one HOTag upon homo-oligomerization of a POI.

**Fig. 2.**
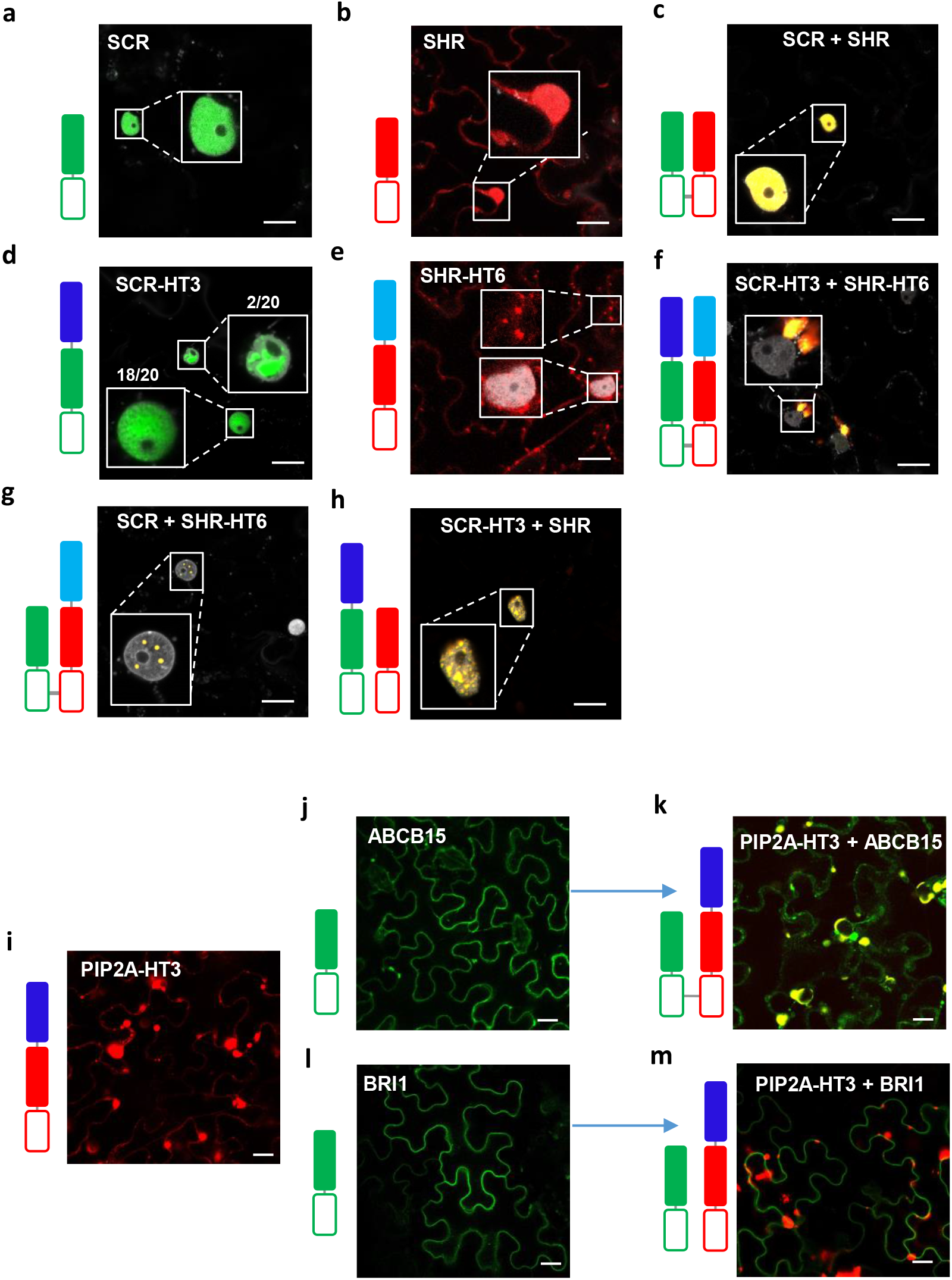
Visualization of transcription factor and membrane protein complexes in plants. **a-h**, Detection of SHR-SCR interactions via HOTag fusions. Localization of SCR-EGFP (**a**), SHR-mScarlet (**b**), co-expression of SCR-EGFP with SHR-mScarlet (**c**), showing uniform expression in the nucleus (**a**,**c**) or nucleus and cytoplasm (**b**). **d**, A small fraction (2 out of 20 nuclei) of SCR-EGFP-HOTag3 (SCR-HT3) formed droplets in the nucleus. **e**, Individually expressed SHR-mScarlet-HOTag6 (SHR-HT6) formed droplets in the cytoplasm. **f**, Co-expression of SCR-EGFP-HOTag3 with SHR-mScarlet-HOTag6 yielded large, dual-colored droplets outside the nucleus. **g-h**, Co-expressing a HOTagged version of one protein with a HOTag-free version of the other partner (SCR-EGFP + SHR-HT6 **(g)** and SCR-HT3 + SHR-mScarlet **(h)**) colocalized in droplets in the nucleus. DAPI staining indicates the position of the nucleus (grey). **i-m**, Detection of PIP2A interactions via HOTag fusions. Localization of individually expressed PIP2A-mScarlet-HOTag3 (PIP2A-HT3) (**i**), ABCB15-GFP (**j**), Co-expression of PIP2A-mScarlet-HOTag3 with ABCB15-EGFP (**k**), BRI1-GFP (**l**) and co-expression of PIP2A-HT3 with BRI1-EGFP (**m**). PIP2A-HT3 always localized in fluorescent droplet-like structures, co-localizing with ABCB15-EGFP but not BRI-EGFP. Individually expressed ABCB15-EGFP and BRI1-EGFP localized at the plasma membrane. All constructs were transiently expressed in *N. benthamiana* leaves, and representative images are presented in the figures. Scale bars = 20 μm.

As expected, the homo- and hetero-oligomerization of SCR and SHR never resulted in phase separation when expressed without HOTags. After combining SCR-EGFP or SHR-mScarlet with SHR-mScarlet-HOTag6 or SCR-EGFP-HOTag3, respectively, we noticed the consistent formation of bright droplets (Fig. 2g-h), reporting SCR-SCR-SHR and SCR-SHR-SHR ternary interactions, respectively. In contrast to the cytoplasmic localization of dual HOTag-labeled SHR-SCR droplets (Fig. 2f) or the single SHR-mScarlet-HOTag6 droplets (Fig. 2e), these “intermediate-valency” droplets were exclusively nuclear, suggesting that in this constellation, SHR-SCR complexes can be imported into the nucleus.

In summary, these data illustrate that SPPIER is a powerful strategy for studying transcription factor complex formation, revealing not only details about homo- and hetero-oligomerization but also about the subcellular localization of the interactions.

### SPPIER can sense interaction between membrane-associated proteins

Integral membrane proteins represent a large proportion of any eukaryotic proteome, often functioning in multiprotein complex assemblies at the center of cellular signaling and endomembrane transport. The higher the number of membrane-spanning domains, the more difficult it is to extract and purify membrane-related complexes, hindering efficient detection of their interaction partners. To evaluate if SPPIER could also be applied for membrane-associated proteins, we selected the water transporting aquaporin PIP2A ^35^, which is known to assemble in homo(tetra-)meric complexes ^36 35 37^. We anticipated that multivalent interactions could not yield *bona fide* liquid-liquid phase separation, as seen for cytoplasmic proteins, due to the membrane insertion of PIP2A.

While PIP2A-mCherry uniformly labeled the plasma membrane (Supplementary Fig. 2), PIP2A-mScarlet-HOTag3 accumulated in large, intracellular fluorescent structures (Fig. 2i), likely derived from PIP2A homo-oligomerization within the endomembrane system. Further, to assess the potential of SPPIER to visualize hetero-oligomerization of membrane-associated proteins, we co-expressed ABCB15-EGFP with PIP2A-mCherry-HOTag3 as both proteins were previously identified in a high-throughput affinity purification experiment as interacting partners ^35^. In the previous experiment (Fig. 2g-h), we established that if one of the partners homo-oligomerizes, the second one does not require HOTag fusion in order to assemble in droplets. Consistently, the co-expressed ABCB15-EGFP co-localized with the PIP2A-mScarlet-HOTag3 in such structures (Fig. 2k). Individually expressed ABCB15-GFP did not show any phase separation (Fig. 2j). This could either corroborate the PIP2A-ABCB15 interaction, or reflect PIP2A-mScarlet-HOTag3 droplet formation pleiotropically interfering with the endomembrane processes, non-specifically accumulating other endomembrane proteins such as ABCB15. To test the latter hypothesis, we co-expressed HOTagged PIP2A with a distinct plasma membrane protein, BRI1. Unlike ABCB15-EGFP, BRI1-EGFP was not recruited to PIP2A droplets, but attained its normal plasma membrane localization (Fig. 2l-m). This excludes the hypothesis that droplet formation interferes pleiotropically with endomembrane trafficking. Taken together, these experiments thus demonstrate that SPPIER can also be applied for detecting PPI between membrane proteins.

### SPPIER sensing of ligand- and phosphorylation-dependent PPI

Subsequently, we evaluated SPPIER for detecting dynamic protein-protein interactions *in planta*. Therefore, we used the strong rapamycin-inducible interaction between the FKBP12 domain and the FKBP12 rapamycin-binding domain of mTOR (FRB) ^38^. Similar to the response in HeLa cells ^20^, FKPB12-EGFP-HOTag3 x FRB-mScarlet-HOTag6 produced rapamycin-induced droplets when transiently co-expressed in *N. benthamiana* leaves (Supplementary Fig. 3 and Supplementary movie 1). Additionally, we demonstrated that a similar Rapamycin-dependent droplet formation could be observed using HOTag4 x HOTag5 and HOTag7 x HOTag2 fusions (Supplementary Fig. 4a-c), thereby providing more flexibility to design more complex experiments using additional HOTags.

To further study the dynamics of droplet formation, we stably co-expressed the FKBP12-EGFP-HOTag3 and FRB-mScarlet-HOTag6 fusion proteins in Arabidopsis cell suspension culture (PSB-D). In line with observations made in HeLa cells ^20^, FKBP12-FRB fluorescent droplets developed within minutes after rapamycin treatment (Fig. 3a and Supplementary movies 2). We quantified the dynamics of droplet formation by calculating the ratio of fluorescent signals present in droplets versus the total fluorescence detected in a cell over time. Using this quantification approach, we observed that within 15 min, about 5-10% of the cellular fluorescence was derived from droplets (Fig. 3a). This demonstrates that the SPPIER strategy enables the detection of inducible interactions in living plant cells, while also showing its utility in plant cell suspension cultures.

**Fig. 3.**
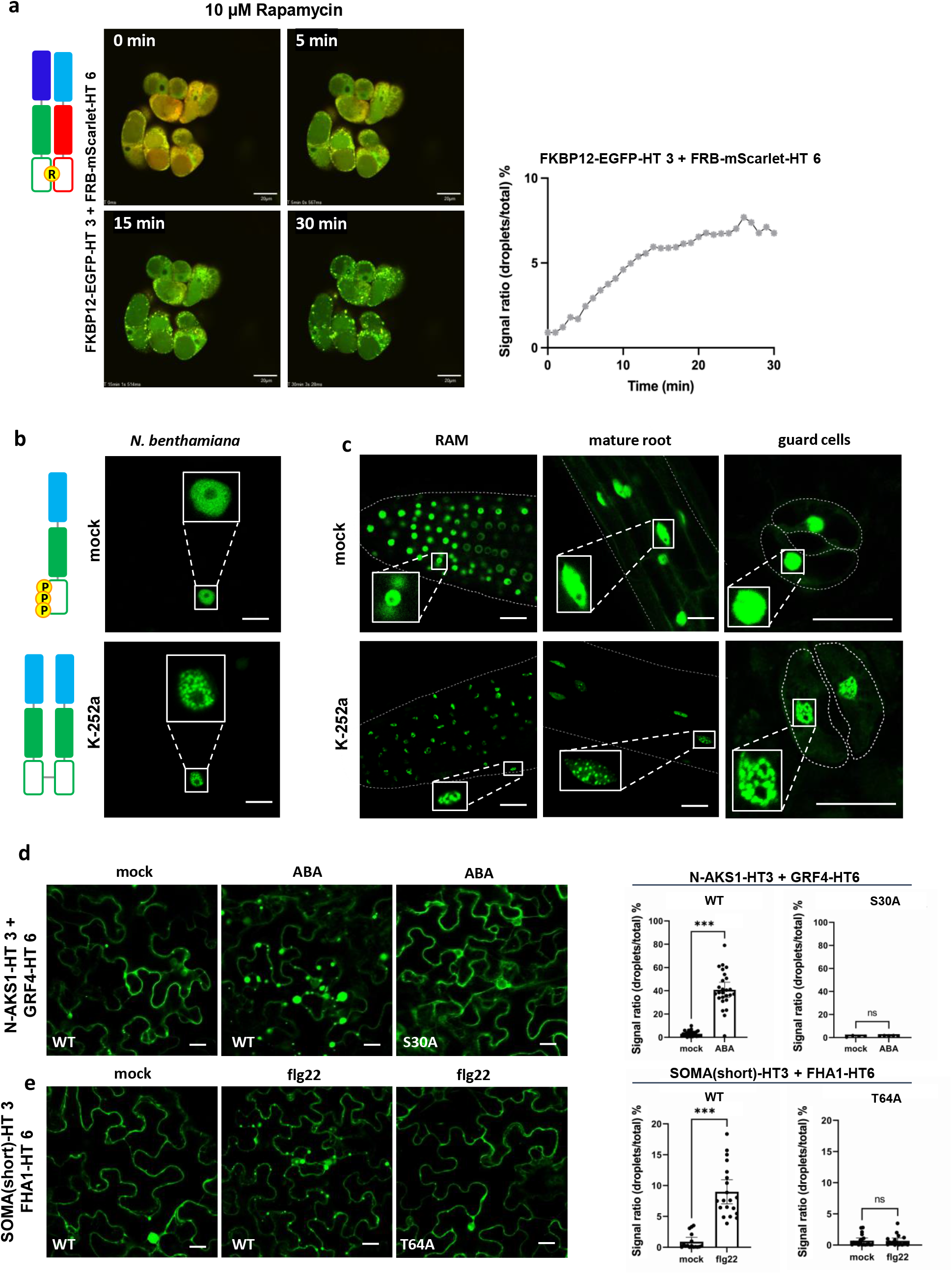
Phase-separation dynamically reports inducible PPI. **a**, Time course of rapamycin-induced (10 μM) droplet formation in Arabidopsis cell suspensions co-expressing FRB-mScarlet-HOTag6 and FKBP12-EGFP-HOTag3 with quantification (right). **b-c**, Fluorescence pattern of AKS1-EGFP-HOTag6 transiently expressed in *N. benthamiana* leaf epidermis **(b)** and in the root apical meristem (RAM), mature root and guard cells of transgenic Arabidopsis seedlings **(c)**. No droplets were observed in steady state conditions. Overnight kinase inhibitor treatment (K-252a) promoted phase separation in the nucleus. **d**, Confocal images and quantification of droplet formation for co-expressed N-AKS1-EGFP-HOTag3 (N-ASK1-HT) with GRF4-EGFP-HOTag6 (GRF4-HT6) in *N. benthamiana* in response to 30 minutes mock (0.1% ethanol) or ABA treatment (50 μM). ABA treatment induced droplet formation in WT, but not in S30A. Average SPARK ratio substantially increases during ABA treatment, compared to the mock treatment. **e**, Confocal images and quantification of droplet formation for co-expressed SOMA(short)-EGFP-HOTag3 (SOMA(short)HT3) + FHA1-EGFP-HOTag6 (FHA1-HT6), treatment with flg22 (1 μM) results in a clear induction of droplet formation. Scale bars = 20 μm. (p<0.05 *, p<0.01 **, p<0.001 ***, Student’s t-test, n>4).

Phosphorylation is an important post-translational modification in many signaling cascades to dynamically regulate the activity of target proteins. In the case of the bHLH transcription factor AKS1, phosphorylation of S284, S288 and S290 in its C-terminus inhibits its homo-dimerization, preventing K^+^ channel expression in guard cells and thus stomatal opening ^39^. In mock-treated *N. benthamiana* epidermal cells, the HOTag-fused full-length AKS1 protein was uniformly expressed in the nucleus (Fig. 3b). Upon treatment with the kinase inhibitor K-252, AKS1-EGFP-HOTag6 appeared in nuclear droplets, suggesting homo-oligomerization of AKS1, consistent with previous *in vitro* results on K-252 induced dimerization of AKS1 ^39^. Similar results were obtained in stably transformed Arabidopsis lines expressing AKS1-EGFP-HOTag6 (Fig. 3c). Interestingly, droplets were not only observed in expected cell types such as guard cells, but also in roots, demonstrating the power of SPPIER for *in planta* visualization of dynamic PPIs at a high signal-to-noise ratio (Fig. 3c).

Interactions between 14-3-3 proteins and their clients are prevalently phosphorylation-dependent ^40 41 42 43^. As an example of this type of interaction, we evaluated our method for the detection of the abscisic acid (ABA)-induced, phosphorylation-dependent interaction between the AKS1 N-terminus (48 amino acids) and the 14-3-3 protein GRF4, which was previously used to develop a Förster resonance energy transfer (FRET)-based kinase activity sensor ^44^. Both components were fused to EGFP followed by different HOTags and co-expressed in *N. benthamiana* leaves. In contrast to the few droplets detected in the mock treatment, droplet formation was strongly induced by ABA treatment (Fig. 3d). When we replaced the phosphorylated Serine in the AKS1 peptide with Alanine (S30A), droplet formation was impaired in the presence of ABA (Fig. 3d). These data demonstrate that SPPIER reliably reports the dynamic phosphorylation-dependent interaction between the AKS1 terminus and GRF4 in living plant cells. Altogether, our analyses illustrate the applicability of SPPIER not only towards stable interactions but also for the detection of dynamic PPIs that are regulated by phosphorylation.

### Analysis of *in planta* phosphorylation through SPARK reporters

As a more versatile alternative to endogenous interaction partners, generic phosphopeptide-binding domains, such as the WW and FHA1 domains, are commonly used in the development of kinase activity readouts. The WW domain specifically recognizes pS-P/pT-P, whereas FHA1 preferentially interacts with pT-X-X-D ^45 46^. The proof-of-principle for using these generic phosphopeptide-binding domains to develop kinase activity reporters based on phase separation was provided in human cell lines where the method was termed SPARK ^20^. To validate the functionality of these general phosphopeptide-binding domains and to a broader extent the usage of SPARK in plants, we first generated HOTagged WW and FHA1 constructs, and co-expressed them transiently in *N. benthamiana* leaves together with compatible phosphopeptides. The WW was combined with an optimized human ERK kinase substrate peptide sequence ^20^ and further co-expressed with the compatible human ERK kinase (hsERK). After treatment with the indirect ERK activator phorbol myristate acetate (PMA), ERK-SPARK droplets were formed ^47^. As expected, both hsERK and PMA were required to enable the induction of ERK-SPARK droplets (Supplementary Fig. 5a-c). Furthermore, droplet formation was lost by mutating the phospho-Threonine in the ERK substrate peptide to Alanine (ERK-SPARK-T48A) (Supplementary Fig. 5d), confirming the phosphorylation-dependency of the droplet formation.

To test the performance of the FHA1 domain in the plant SPARK approach, HOTagged FHA1 was combined with a 90 amino acid peptide sequence that was used in the plant FRET-based Mitogen-Activated Protein Kinase (MAPK) activity reporter (sensor of MAPK activity; SOMA) ^48^. Unlike the FRET counterpart, the SPARK reporter with the original SOMA peptide displayed constitutive activity in *N. benthamiana*, suggesting the presence of additional phosphorylation sites in the original SOMA peptide that were previously not detected ^48^ (Supplementary Fig. 6a). We could abolish this non-specific activity by trimming the original 90 amino acid substrate peptide down to 15 amino acids (Fig. 3e and Supplementary Fig. 6b), which is in line with the typical range of substrate sequences that are used to develop highly specific kinase activity sensors (8-18 amino acids) ^20^. The Serine (S71) in this peptide was replaced by Alanine to exclude its off-target phosphorylation. Using the trimmed peptide, the SOMA-SPARK produced very few droplets in control conditions, while showing a very clear droplet induction upon flagellin epitope (flg22) treatment (Fig. 3e). This response was specific to T64 phosphorylation, as droplet formation was impaired when this Threonine was mutated to Alanine (SOMA-SPARK-T64A) (Fig. 3e). Jointly, these data demonstrate that WW and FHA1 can be used for developing SPARKs in plants for detecting the phosphorylation status of a phosphopeptide of interest, in the context of an endogenous signaling cascade, as well as for determining kinase-substrate relationships.

### Development of a SPARK-based SnRK1 activity sensor

Having established that the SPARK strategy works for *in vivo* kinase activity detection in plants (Fig. 3e and Supplementary Fig. 5), we deployed the SYMPL vectors to generate a novel kinase activity reporter for SnRK1. SnRK1 protein kinase is the plant ortholog of the yeast SNF1 and the mammalian AMPK kinases and functions as a metabolic sensor that is essential for survival under energy deprivation ^49^. Because of this central role in metabolism, SnRK1 plays a crucial role in stress responses, while also orchestrating plant development. Currently, however, no real-time *in vivo* SnRK1 kinase activity reporters are available, hindering efficient evaluation of the dynamics of SnRK1 activity in a spatio-temporal context under different environmental conditions. Based on the strong conservation of the AMPK/SNF1/SnRK1 phosphorylation recognition motif ^22^, we selected the synthetic AMPK Substrate Peptide (ASP) as a likely substrate sequence for SnRK1. This ASP was previously used in combination with the FHA1 domain for the generation of *in vivo* ^50^ or *in vitro* ^51^ mammalian AMPK reporters. The phosphorylated Threonine (T10) in the ASP is followed by an aspartic acid (D) on position +3, making ASP thus highly compatible for binding to the FHA1 domain upon phosphorylation by SnRK1. Therefore, we designed a novel plant SnRK1 kinase reporter combining ASP-EGFP-HOTag3 and FHA1-mCherry-HOTag6 as components for SPARK (Fig. 4a).

**Fig. 4.**
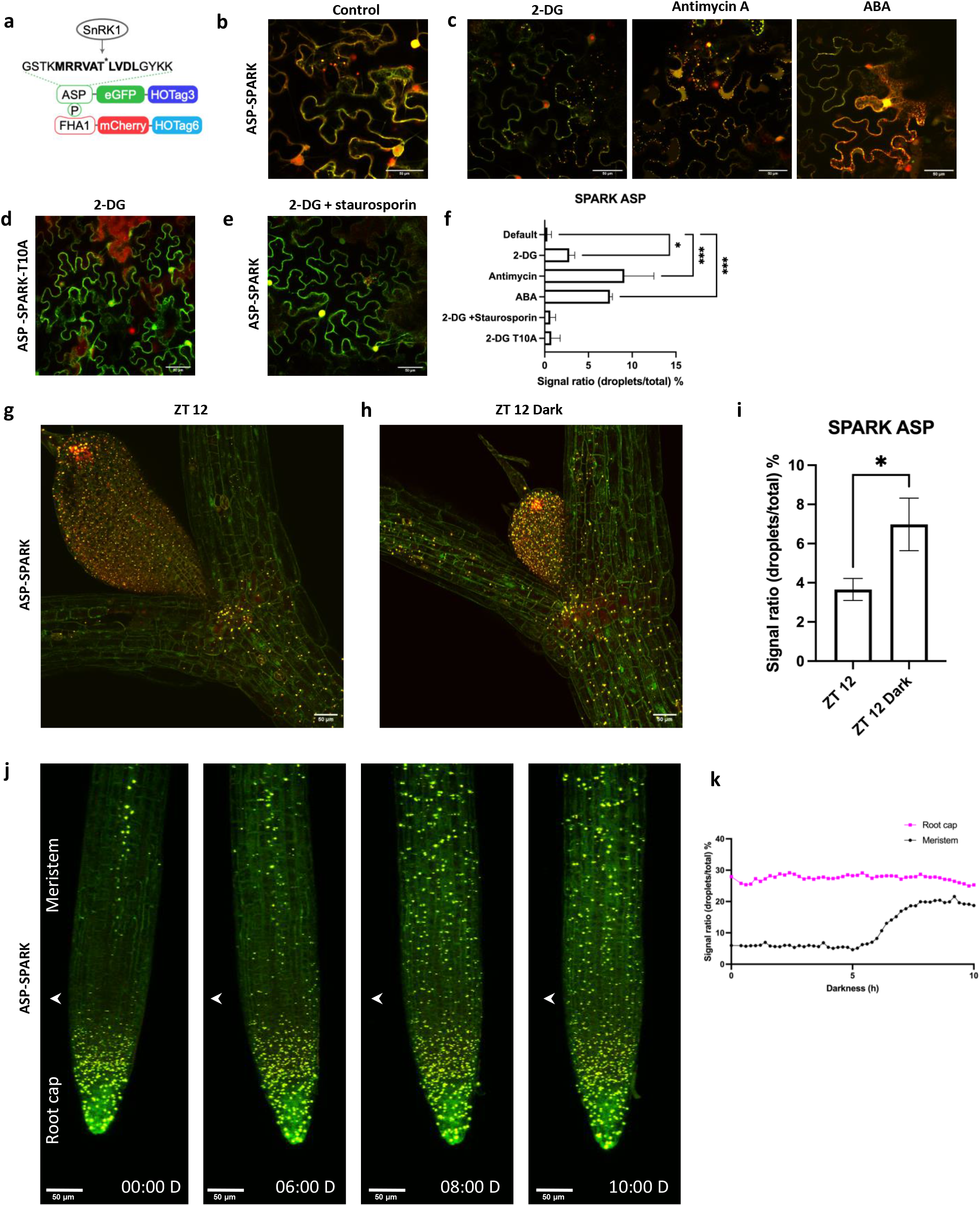
Development of an ASP-SPARK sensor for *in planta* monitoring of SnRK1 kinase dynamics. **a**, Schematic overview of ASP-SPARK, consisting of overexpression cassettes for the ASP-AMPK consensus motif fused to EGFP and HOTag3 (ASP-HT3), and FHA1 fused to mCherry and HOTag6 (FHA1-HT6). SnRK1-mediated ASP-HT3 phosphorylation induces interaction with FHA1-HT6. **b-c**, Transiently co-expressed ASP-HT3 with FHA1-HT6 (ASP-SPARK) in *N. benthamiana* leaves **b**, untreated, or **c**, treated for 30 min with 10 mM 2-DG, 20 μM antimycin A or 5 μM ABA. A substantial droplet formation was induced by the treatments. **d**, Droplet formation of the mutated ASP-SPARK-T10A was insensitive to 2-DG. Scale bars =20 μm. **e**, Pre-incubation with 10 μM staurosporine (2 hours) prevented 2-DG induced droplet formation. Scale bars = 20 μm. **f**, Quantification of the SPARK ratio in the infiltrated *N. benthamiana* leaves after treatments described in a-d (n=3). **g**, Droplet formation in ASP-SPARK seedlings (8DAG), 12h after onset of photoperiod (ZT 12). **h**, ASP-SPARK seedlings kept in darkness for 12 hours after onset of photoperiod (ZT 12 Dark). **i**, Quantification of the SPARK droplet formation from **g** and **h** (n=4). SPARK ratios were calculated for the whole image. **j**, Time lapse of an ASP-SPARK root after transfer to sugar free medium and kept in the dark to activate the SnRK1 low-energy responses. **k**, Quantification of the phase-separation response in the root tip of the experiment in (j). Phase separation was measured from the top of the image down to the arrowhead (meristem), and below the arrowhead (root cap).

To explore if the ASP peptide could indeed serve as a substrate for the Arabidopsis SnRK1 kinase, we first used SPPIER to test the interaction between the ASP peptide and SnRK1α1, the catalytic subunit of the heterotrimeric SnRK1 complex, in *N. benthamiana* leaves. In the control experiment with individually expressed SnRK1α1-mScarlet-HOTag3, fluorescent droplets appeared (Supplementary Fig. 7a), in agreement with the known homo-multimerization of the human AMPK and yeast SNF1 kinase complexes ^52 53 54^. This indicates that also the plant SnRK1 kinases can form higher-order oligomeric complexes, in accordance with the observed dimerization of the regulatory SnRK1βɣ subunit in maize ^55^. On the other hand, in the control experiment with the single ASP-EGFP-HOTag2 component, no droplets were observed (Supplementary Fig. 7a). Importantly, co-expression of both ASP-EGFP-HOTag2 and SnRK1α1-mCherry-HOTag3 induced dual-colored fluorescent droplets (Supplementary Fig. 7a), proving direct interaction between ASP and SnRK1α1. This interaction was not abolished by mutating the phosphorylatable Threonine residue in the ASP peptide to Alanine (Supplementary Fig. 7b), illustrating that the Threonine residue of ASP is not required for the kinase-substrate interaction of ASP and SnRK1α1. To further explore the specificity of this interaction, we co-expressed ASP-EGFP-HOTag2 with SnRK2.2-mScarlet-HOTag3, a member of the closely related ABA-specific SnRK2 kinase family ^56^ (Supplementary Fig. 7a). The absence of dual-colored droplets in this SPPIER combination suggests that ASP does not interact with SnRK2 kinases, indicating that ASP interacts specifically with SnRK1α1.

After obtaining evidence for the interaction between ASP and SnRK1α1, we next evaluated the performance of the ASP-SPARK reporter as SnRK1 kinase activity sensor via transient expression in *N. benthamiana* leaves and stable expression in Arabidopsis cell suspension cultures (PSB-D). Under default, non-stressed conditions, almost no ASP-SPARK droplets were observed (Fig. 4b and Supplementary Fig. 8a). In contrast, inhibition of glycolysis with 2-deoxyglucose (2-DG) ^49^ or interference with the electron transport chain by Antimycin A ^57^, two agents known to stimulate SnRK1 through activation of low energy responses ^58 59^, significantly induced ASP-SPARK droplet formation (Fig. 4c,f and Supplementary Fig. 8a). In addition, droplet formation was also triggered by the stress hormone ABA, consistent with earlier findings of ABA-dependent SnRK1 activation ^60 61^ (Fig. 4c,f). The ASP-SPARK response was dependent on the phosphorylation of its Threonine residue, as droplet formation was impaired for the mutated ASP peptide (ASP-T10A) (Fig. 4d,f and Supplementary Fig. 8b). This phosphorylation dependency was further corroborated through pre-incubation of the leaves with staurosporine, a general kinase inhibitor, which also prevented droplet formation (Fig. 4e-f).

Taking into account the specific interaction between ASP and SnRK1, as well as the responsiveness of the ASP-SPARK reporter towards activators of SnRK1, we finally generated transgenic *Arabidopsis thaliana* lines constitutively expressing both components of the ASP-SPARK reporter. Given its ubiquitous and high expression in Arabidopsis, we selected the ASPARTIC PROTEASE A1 (APA1, AT1G11910) promoter, which was recently used in other *in vivo* plant sensors ^62^, to drive the expression of the ASP-SPARK reporter. Seedlings overexpressing ASP-SPARK construct were indistinguishable from wild-type plants and did not show any perturbations in seedling growth. Furthermore, ASP-SPARK plants did not show an altered response to energy deprivation evoked by submergence and darkness, suggesting that its constitutive expression does not interfere with endogenous SnRK1 functioning or plant development (Supplementary Fig. 9a-c) ^63 64^. Analysis of the fluorescence signals in the shoot tissue of the ASP-SPARK seedlings (8 DAG) revealed that during the day (12 hours after the onset of the photoperiod (ZT 12)), droplets were primarily confined to the developing leaves primordia (Fig. 4g). Likewise to the ASP-SPARK results obtained in *N. benthamiana*, the dual-colored ASP-SPARK droplets were sensitive to staurosporine treatment, while also being dependent on the phosphorylatable ASP-T10 residue (Supplementary Fig. 8c-d), confirming their phosphorylation-dependency *in planta*. To explore the responsiveness to low energy, we exposed ASP-SPARK seedlings to an extended night treatment in the absence of an exogenous sugar source, a condition known to stimulate SnRK1 activity ^65 66^. After a 20h period of darkness (seedlings were kept in the dark for 12 hours after the onset of photoperiod (ZT 12 dark)), droplet formation increased both in the developing leaves as well as in the cotyledonary nodes, illustrating that the ASP-SPARK line monitors low energy induced kinase activities in entire organs at cellular resolution (Fig. 4h-i). Recently, it was shown that dark treatment also impacts SnRK1-mediated signaling in the root ^67^. Therefore, we explored the spatio-temporal characteristics of the root response to unexpected darkness using the ASP-SPARK reporter over a 20h time period using a tilted confocal microscope and automated tracking of the growing root tip. At the onset of the treatment (ZT 12), ASP-SPARK droplets were mainly restricted to the most rootward part of the meristem and root cap. Furthermore, droplets appeared in some cells of the vascular bundle as they reached the elongation zone. After more than 6h of darkness, droplets appeared across the entire root meristem, indicating induced SnRK1 activity in response to the dark treatment (Fig. 4j-k and Supplementary Movie 3). In summary, these results demonstrate that the developed ASP-SPARK reporter represents a powerful novel tool for real-time exploration of dynamic SnRK1 signaling at an unprecedented spatio-temporal resolution under different environmental conditions.

## Discussion

Phase-separation due to multivalent interactions is widely spread in biology, serving to spatially organize and biochemically regulate cellular processes. In this work, we showcase that phase separation by synthetic multivalency can serve as a versatile approach to monitor PPIs and phosphorylation events *in planta*. In a remarkably simple design, installing synthetic multivalency between two monovalently interacting proteins only requires their fusion to homo-oligomeric tags. Co-expression of both components is then sufficient to trigger liquid-liquid phase separation of the complex, enabling sensitive detection at a high signal-to-noise ratio via fluorescent markers included in the fusion proteins. The proof of principle for the use of paired HOTags for detecting PPI and kinase activities in the cytoplasm was previously provided in animal cell lines ^20 19^, but remains poorly explored. Here, we generated a versatile toolbox of Golden Gate and Gateway compatible modules for efficient cloning of HOTagged protein fusions, and used them to establish the technology for plant-based systems.

In addition to demonstrating its utility for analyzing interactions among cytoplasmic proteins in tobacco leaves, Arabidopsis cell suspension cultures and seedlings, we found that the synthetic multivalency also allows the detection of interactions between nuclear, as well as integral membrane proteins. The technology also easily detects homo-oligomerization, which is a feature that could complexify the dissection of the different interactions in a complex. To identify homo-oligomerization it was sufficient to express individual HOTagged constructs of PIP2A, SHR, SCR, or SnRK1α1. Additionally, co-expression of a single HOTagged homo-oligomerizing protein and a non-HOTagged interactor yielded dual-labeled droplets. This indicates that the induced “intermediate-valency” due to homo-oligomerization of one interaction partner is sufficient to trigger phase separation of a complex that could still recruit additional interacting partners. Unlike the previous reports ^20 19^, we were able here to upgrade the sensitivity and the detection capacity of this approach by simply employing distinct fluorescent proteins. This enabled us not only to visualize higher-order interactions (TPLATE-TASH3-LOLITA) but also to infer two readouts from single experiments such as SHR homo-oligomerization and SHR-SCR interaction, SCR homo-oligomerization and its interaction with SHR, and PIP2A homo-oligomerization next to PIP2A-ABCB15 interaction. The simultaneous reporting of two interactions based on only two constructs is unprecedented for fluorescence-based reporters.

A high concentration of the multivalent interactors is required for reaching the critical threshold for phase separation ^18^. Consequently, all components involved in the multivalent interaction need to be present at sufficient levels. On the one hand, this calls for caution when interpreting phase separation-based reporters in a developmental context, as the sensitivity of the sensor can be modified if the expression levels and protein concentration of either component are not stable in the cells or tissues of interest. On the other hand, this characteristic of the technology opens new venues for studying higher-order interactions, as the expression levels of endogenously expressed interaction partners are expected to be too low to trigger significant phase separation. Here, this is shown by the inability of endogenously expressed TASH3 to connect sufficient HOTagged TPLATE and LOLITA to obtain phase separation in *N. benthamiana*. The critical concentration of this ternary complex was only achieved upon co-overexpression of TASH3-TagBFP. This illustrates how phase separation permits robust dissection of ternary interactions. One could further expand the number of possible interactions detected by the additional co-expression of other, non-HOTagged, fluorescently labeled interaction partners. In this case, one expects the additional co-localization of the new interaction partner into the droplets, alike the recruitment of non-HOTagged proteins to the dual-readout with homo-oligomer droplets mentioned above.

One of the endogenous functions of phase separation in biology is to control the localization and thus the activity of its components as seen for ARF transcription factors ^16^. Consequently, it is to be expected that the proteins recruited to phase-separated droplets do not attain their normal localization. However, we found that phase separation does not completely eliminate all subcellular details of interaction dynamics, revealing that SHR-SCR and SHR-SHR interaction are established in the cytoplasm. Also, the plasma membrane protein PIP2A accumulated in intracellular structures. Importantly, this did not cause pleiotropic endomembrane trafficking defects, as BRI1-EGFP could still reach the plasma membrane.

From our analyses, it is clear that the HOTags allow the *in planta* detection of strong direct interactions between two proteins as well as ligand-induced interactions (Fig. 1-3 and Supplementary Fig. 3-4). The critical need for a *bona fide* interaction to establish multivalency indicates that this strategy is less prone to false positives related to strong overexpression of the sensors, compared to proximity-based technologies such as Bimolecular fluorescence complementation (BiFC) ^2 3^.

Importantly, our strategy also enables the *in planta* detection of weak electrostatic interactions between a phospho-binding domain and a phosphorylated peptide with very high sensitivity and specificity. Different phospho-binding domains could be used to sense the phosphorylation of two distinct consensus sequences (pS-P/pT-P for WW and pT-X-X-D for FHA1). This could impose restrictions on the sequence of phosphosites that could be detected. However, many kinases tolerate modifications of the substrate sequence given that they do not disrupt the recognition sequence. For example, a given candidate sequence could be modified to contain a D at the +3 position to match the FHA1 consensus sequence without disturbing the substrate recognition site. The high sensitivity of the SPARK calls for a need for short substrate sequences that are devoid of secondary phosphorylation sites. We obtained specific responses using substrate sequences between 15 and 48 amino acids long, whereas the 90 amino acid SOMA substrate contained additional phosphorylation sites that were sensed by the FHA1. The minimal substrate sequence is, however, not always sufficient for recognition by some kinases. For example substrate binding by MAPKs depends on a docking site outside the actual phosphorylation domain ^68^.

In its simplest application, SPARKs can be used to validate phosphorylation events that were detected via phosphoproteomics. We also demonstrated the use of SPARKs to evaluate the kinase-substrate relationships for human ERK and an ERK substrate via an *in planta* kinase assay, where the kinase is co-expressed with a substrate-based SPARK (Supplementary Fig. 5). Lastly, we applied all principles mentioned above to develop a reporter for SnRK1, allowing us to detect with unprecedented resolution the induction of SnRK1 kinase activity in a vertically growing root with cellular resolution.

The interaction and recruitment of interactors to phase-separated droplets might have dominant-negative effects on the signaling cascade of interest. As we were able to obtain normal-looking, stable lines expressing ASP-SPARK, we hypothesize that such dominant-negative effects are manageable. Besides, the interpretation of the SPARK readout in different developmental contexts should be done with caution, as different cell types and cell volumes could impact the protein concentration and thus the cellular sensitivity of the SPARK reporter.

To date, studying a single phosphorylation event via mass spectrometry, phospho-specific antibodies or even FRET-based sensors represents a technical challenge to researchers. Considering minimal optimization, the ease of cloning and high sensitivity using basic confocal microscopy, our SYMPL toolbox now makes it possible to rapidly develop a sensor for specific phosphorylation events in plants. Therefore, we envision that SPARK and derivatives thereof will revolutionize the study of phosphorylation-based signaling in plants. Furthermore, expansion of the SYMPL toolbox can also open the door for detecting and monitoring a wide assortment of post-translational modifications, such as ubiquitination or acetylation, by swapping the FHA1/WW/14-3-3 phospho-binding domain with a ubiquitin or acetyl-Lysine binding domain ^69 70 71^. Additionally, this approach can be adapted to study other types of biomacromolecule interactions involving for instance protein-RNA recognition ^12 72^. *In vitro* studies were even able to specifically visualize phase-separated protein-methylated DNA interaction ^73^.

To conclude, we illustrated the easy design principles, versatility and robust output of SPPIER and SPARK sensors for the *in planta* study of basic and complex PPIs and even dynamic phosphorylation changes, with unprecedented sensitivity. We provide the SYMPL toolbox for its wide implementation and further development as a catalyst for the routine, as well as more complex studies of PPIs and post-translational modifications in plants.

## Methods

### Cloning

EGFP-HOTag3 and EGFP-HOTag6 were amplified by PCR from pcDNA3-ERK-SPARK ^20^ and inserted in pDONRP2RP3 via Gateway BP reaction for C-terminal fusion using MultiSite Gateway-based cloning. The EGFP sequences were then replaced by mScarlet via Gibson assembly to create mScarlet-HOTag3 and mScarlet-HOTag6. In order to test the interaction of two given proteins, their coding sequences (CDS) were amplified by PCR and inserted in pDONR221 via BP reaction. Next, the first CDS was fused to EGFP-HOTagX and the second to mScarlet-HOTagY by performing a MultiSite Gateway LR reaction both under the control of the CaMV 35S promoter.

SPARK expression vectors were created via cut-ligation ^74^. Substrate peptide sequences were ordered as oligonucleotides (IDT®). WW, FHA1 and 14-3-3 (GRF4) sequences were PCR-amplified, respectively, from pcDNA3-ERK-SPARK ^20^, pET-SOMA ^48^ and Arabidopsis Col-0 cDNA by using the Q5 DNA polymerase (NEB®). The resulting PCR fragments were redeemed via column purification. The resulting DNA fragments were cloned into the PGGB000 GreenGate entry vectors via cut-ligation and verified via Sanger sequencing. LinkerI-EGFP was PCR amplified from the pcDNA3-FRB-EGFP-HOTag6 ^20^ vector and was cloned into the PGGC000 Greengate entry vector. The HOTag DNA sequences were ordered as gBlocks® from IDT® and cloned into the pGGD000 Greengate entry vector via cut-ligation. General Greengate building blocks (pGGA-pCAMV35S-B, PGGA-pUBI10-B, pGGC-EGFP-D, pGGC-mCherry-D, pGGC-PmScarlet-I-D, pGGE-G7t-F, pGGE-OCSt-F, pGG-E-35St-F, pGGF-KanR-G, PGGF-tNOS-G, PGGF-sacB-G, pGGF-linkerII-G) and expression vectors (pK-G-U1-ccdB-A-U9, pP-A-ccdB-G, pK-A-ccdB-G) were obtained from the PSB plasmid repository (https://gatewayvectors.vib.be). The PGG-A-pAT1G11910 (pAPA1)-B vector was kindly provided by prof. Karin Schumacher from the Center for Organismal Studies (COS) from the University of Heidelberg ^62^.

For the *in vivo* kinase assay, the *hsERK* coding sequence (Addgene #82145) was introduced in the pHCTAP destination vector ^75^ via MultiSite Gateway cloning. pHCTAP contains a separate RFP expression cassette to detect the cells that contain this vector.

The ASP-SPARK and FRB-FKBP12 constructs were cloned via Golden Gibson. First, each subunit was cloned into a Gibson entry vector via Golden Gate assembly. I-SceI-digestion for 2h at 37°C was then used to linearize the entry and destination vectors (pGGIB-U1-subunit I-U2, pGGIB-U2-subunit II-U3, pGGIB-linkerIII-U3, pK U1-A-ccdB-G-U9). 5 μl NEBuilder HiFi® was added to 5 uL of the digestion and incubated at 50°C for 1h. 1 uL of the reaction mixtures was transformed into 50 μl of electrocompetent DH5a *E. coli* cells and plated out on LB selective medium containing the 75 μg/ml Spectinomycin. Vectors were checked via PCR and verified by Sanger sequencing and subsequently transformed into Agrobacterium C58C1 RifR (pMP90) via electroporation.

### Plant growth conditions and transformation

Tobacco plants were grown for 4 weeks in the greenhouse (16h light at 26°C, 8h dark at 21°C). Leaves were infiltrated with Agrobacterium C58C1 RifR (pMP90) cells at an OD600 of 0.5 for each construct, dissolved in infiltration buffer (100 μM Acetosyringone, 10 mM MgCl2 10 mM MES (pH5.7)) using a 1 ml syringe. A p19 antiviral suppressor gene was co-infiltrated to suppress viral defense responses ^76^. 2-3 days after transformation leaf excisions were imaged via confocal microscopy. ABA, flg22, 2-Deoxyglucose, Antimycin A, staurosporine and rapamycin were dissolved in the same infiltration buffer (lacking Acetosyringone) at the appropriate concentrations (respectively 50 μM, 1 μM, 20 mM, 20 μM, 5 μM and 10 μM) and infiltrated into the transformed leaf area with a 1 ml syringe.

Arabidopsis PSB-D cultures were transformed via co-cultivation with *Agrobacterium tumefaciens* and weekly maintained as earlier described ^77^. Treatments to induce SnRk1 activity were applied 7d after renewal by spiking the MSMO medium with 2-DG, Antimycin A and ABA at the appropriate concentrations (respectively 50 mM, 20 μM and 10 μM). Treated cell culture samples were gently agitated on an orbital shaker for 8h until imaging was performed.

To generate stable plant lines, *Arabidopsis thaliana* Col-0 plants were transformed with the AKS1-EGFP-HOTag6, and ASP-SPARK constructs via floral dip. Transformed plants were selected on ½ Murashige & Skoog medium (½ MS) containing 50 μg/L Kanamycin. Plants were grown under long-day conditions (16h light at 21°C, 8h dark at 18°C). For extended darkness treatment plants were grown on plates with ½ MS medium containing 1% sucrose under long-day conditions, 7 DAG plants were transferred to ½ MS plates without sucrose and kept in dark for 20h before imaging. For staurosporine treatment, plants were transferred on 7 DAG towards liquid ½ MS medium without sucrose containing 10 μM staurosporine and incubated overnight.

For the complete energy deprivation assay, ASP-SPARK and WT Arabidopsis plants were grown under long-day conditions (16h light at 21°C, 8h dark at 18°C). 6 weeks after germination, plants were submerged and/or kept in complete darkness. After 36h plants were transferred back to the greenhouse and further grown under long-day. 7 days later, survival was scored.

### Imaging

Confocal images were taken on a Zeiss LSM710 inverted confocal microscope or an Olympus FluoView1000 (FV1000 ASW) laser point scanning microscope, equipped with a Zeiss C-Apochromat 40x/1.2 water-corrected M27 objective or an Olympus Plan-Apochromat 20x/0.75 dry objective lens respectively. For the nuclear proteins, images were acquired with a Zeiss LSM710 using a 63x water-corrected M27 objective (NA 1.2).

EGFP was excited with an Argon laser at 488 nm and emission was acquired in the range from 500 to 530 nm. mCherry/mScarlet was excited with a solid-state 559 nm laser and emission light was acquired between 600 and 630 nm. The pinhole was set to 1 Airy unit and the images were acquired at 1024 × 1024 pixels resolution for the Zeiss and 512 × 512 pixels resolution for the Olympus with a bit depth of 12 (Olympus) or 16 (Zeiss) using sequential scanning. For SPARK experiments, the first 2-3 layers of cells alongside the cut leaf were omitted since wounding tends to induce non-specific droplet formation, especially with SOMA. All samples were mounted between slide and coverslip and imaging of dual labeled samples was done using line sequential mode.

Live imaging of the growing root tip was executed with the inverted Zeiss LSM900 KMAT confocal microscope equipped with a Zeiss Plan-Apochromat 20x/0.8 dry objective. Eight-day-old Arabidopsis seedlings were placed on a house-made two-part chambered cover glass (adapted from https://cellgrowth-lab.weebly.com/3d-prints.html) and covered with a block of ½ MS without sucrose and 1% agar. For GFP detection a 488 nm Diode laser 10mW and a GaASP-PMT detector (490-565 nm) were used. mCherry was excited with a 561 nm Diode laser (SHG) 10mW, and emission was detected with a GaASP-PMT detector (578-696 nm). Images were acquired at 512×512 pixel resolution with 8-bit depth and a time interval of 720 s. Tracking of the growing root tip was done using a Tip track Matlab script described in ^78^. Confocal images were processed by using the Zeiss Zen 2.3 Lite software package and the FIJI software package ^79^.

### Quantification of the SPARK signal

Quantification of SPARK ratios was done by using FIJI software as follows:

1. Open the image with imageJ.
2. Select the EGFP-channel.
3. Quantify the total signal.
  a. Analyze>measure (the chosen parameters: Area, Mean, Min, Max).
  b. Multiply the mean signal intensity by the total area.
4. Quantify the droplet signal.
  a. Image>Adjust>Threshold>Shanbhag.
  b. Adjust the threshold
  c. If needed, manually exclude the nucleus (unless the phosphorylation is studied in the nucleus) and the non-specific oversaturated signal. Select the area you need to exclude and press “delete”.
  d. Analyze>measure.
  e. Multiply the mean signal intensity by the total area of the droplets.
5. Divide the droplet signal by the total signal of the entire image.

### Statistical analysis

All statistical analyses were performed in Graphpad Prism 9. All plotted values are mean +/-SEM. Unless stated otherwise, p-values were calculated using one-way ANOVA followed by an uncorrected Fisher’s LSD test (p<0.05 *, p<0.01 **, p<0.001 ***). Sample sizes are mentioned in the captions of the according graphs.

## Acknowledgments

This work was funded by a FWO-SB fellowship to W.S. (1S20521N), a FWO Research Grant to S.V. (G002220N) and G.D.J (G011720N), a European Research Council (ERC) to D.V.D (T-Rex project number 682436), and a FWO grant (G.0C76.18N), the LABEX CEMPI (ANR-11-LABX-0007) and I-SITE ULNE (ANR-16-IDEX-0004) to F.B.R.

## Author contributions

A.S. W.S., J.V.L and S.V. wrote the manuscript with the input of all co-authors.

A.S, W.S., B.C., F.B.R., T.B., G.D.J., J.V.L. and S.V. conceived the research.

A.S., W.S., X.K., A.F., and B.C. executed the experiments.

F.B.R., D.G., G.J., T.B. J.V.L. and S.V. supervised the research.

A.G., E.M. and D.V.D. assisted during confocal imaging and image analysis.

## Notes

### Competing Interest Statement

The authors have declared no competing interest.

