## Supplementary Figures for "Phase separation-based visualization of protein-protein interactions and kinase activities in plants"

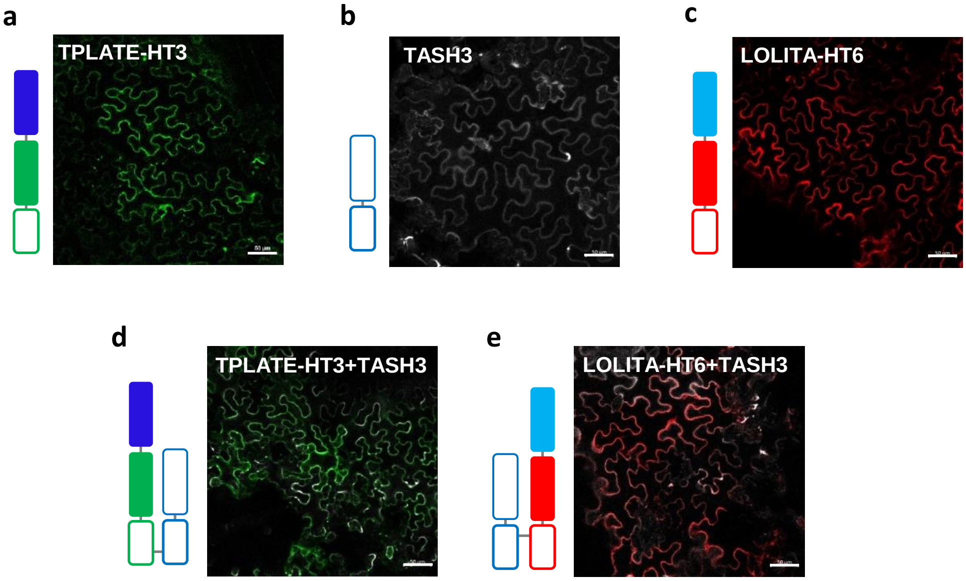


**Supplementary Fig. 1 l Controls for ternary interactions among TPC components.** Representative confocal images of *N. benthamiana* leaves transiently expressing different combinations of TPC proteins. **a,** TPLATE-EGFP-HOTag3 (TPLATE-HT3), **b,** TASH3-TagBFP2 (TASH3), **c,** LOLITA-mScarlet-HOTag6 (LOLITA-HT6), **d,** TPLATE-HT3 + TASH3, **e,** LOLITA-HT6 + TASH3. Neither the individually-expressed proteins **(a-c)** nor co-expression of pairs of the TPLATE/LOLITA/TASH3 trimer **(d-e)** yielded phase-separated puncta. Scale bars = 50 µm.

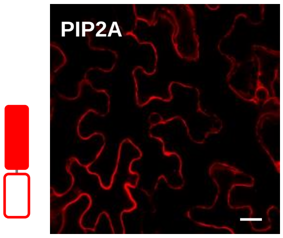


**Supplementary Fig. 2 l The HOTag-free plasma membrane aquaporin PIP2A does not form droplets.** Representative confocal images of *N. benthamiana* leaves transiently expressing PIP2A-mCherry (PIP2A). Scale bars = 20 µm.

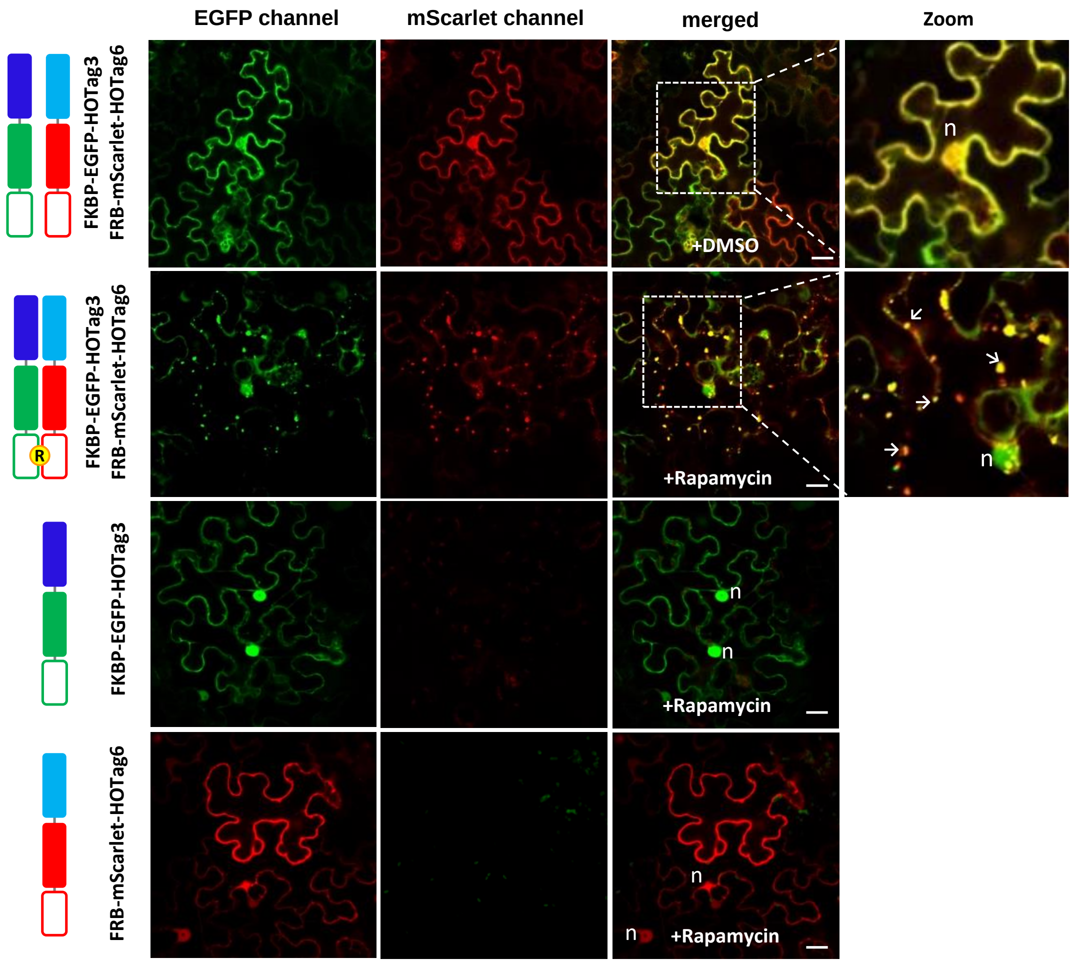


**Supplementary Fig. 3 l HOTags can be used to monitor the rapamycin inducible FRB-FKBP12 protein-protein interaction in plants.** Rapamycin treatment (10 µM for at least 10 min) of *N. benthamiana* cells transiently expressing FKBP12-EGFP-HOTag3 and/or FRB-mScarlet-HoTag6. Arrows indicate droplets. n=nucleus. Scale bars = 20 µm.

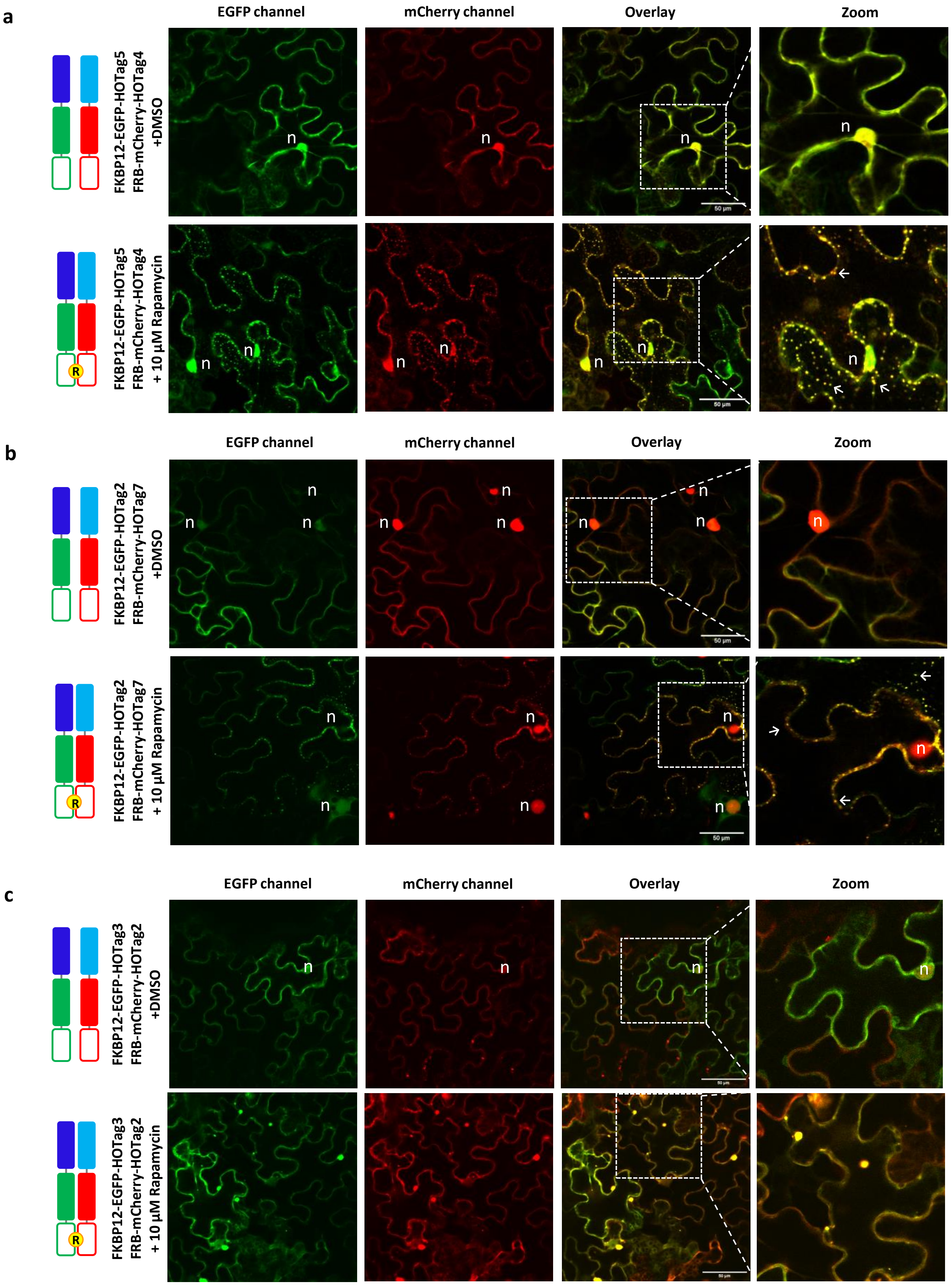


**Supplementary Fig. 4 l Different combinations of HOTag report rapamycin-inducible PPIs. a,** Confocal images of *N. benthamiana* leaves transiently co-expressing FKBP12-EGFP-HoTag4 and Frb-mScarlet-HoTag5 **b**, or FKBP12-EGFP-HoTag7 and FRB-mScarlet-HoTag2 **c,** or FKBP12-EGFP-HoTag3 and FRB-mScarlet-HoTag2. Each combination was treated with DMSO or 10 µM Rapamycin and imaged after 45 min. The rapamycin treatment induced the formation of dual colored droplets. Scale bars = 50 µm. Droplets are indicated by arrows, nuclei by n.

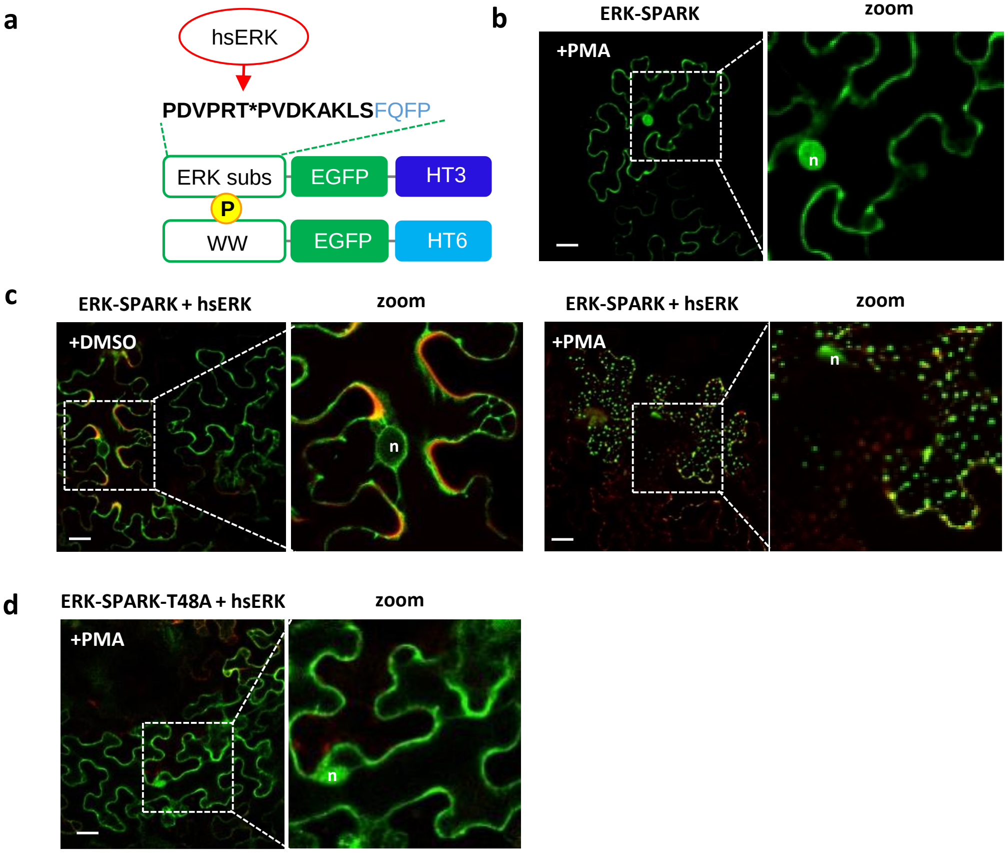


**Supplementary Fig. 5 l SPARK allows monitoring kinase activity dynamics. a,** Scheme of ERK-SPARK, containing hsERK's substrate sequence (bold) and docking sequence (blue) **b**, Transiently expressed ERK-SPARK (contains hsERK's target, cdc25-derived peptide) without co-infiltration of hsERK (hsMAPK1) in *N. benthamiana* leaves, treated with 1 µM PMA. **c.** Transiently expressed ERK-SPARK with co-infiltration of hsERK in *N. benthamiana* leaves, treated with DMSO (mock) and 1 µM PMA for 30 minutes. **d,** ERK-SPARK-T48A which is missing the phosphorylatable Threonine did not produce droplets in any condition. n=nucleus. Scale bars = 20 µm.

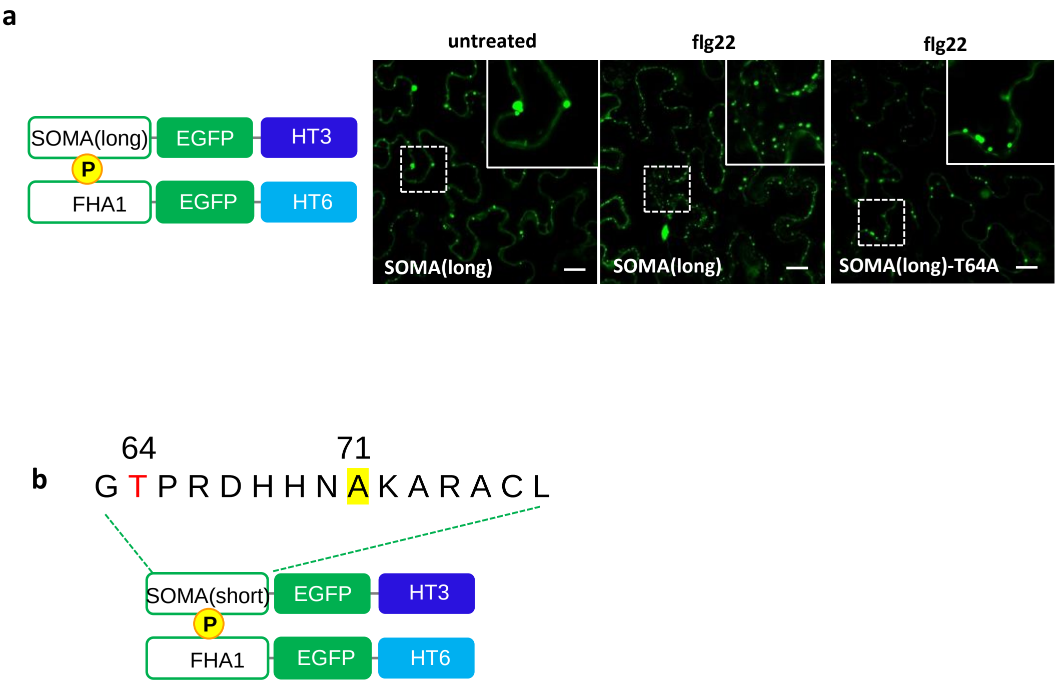


**Supplementary Fig. 6 l The long version of SOMA peptide (90 AA) shows high background signal and non-specific SPARK.** **a,** Scheme of SOMA(long)-SPARK (co-overexpression of FHA1-EGFP-HOTag6 and SOMA(long)-EGFP-HOTag3). SOMA(long)-SPARK formed droplets in both treated (1 µM flg22 for 30 min) and non-treated leaves in WT as in T64A. Insets display details of droplet formation. **b,** The 15 amino acid sequence used in the optimized SOMA(short). The phosphorylatable T64 is shown in red. The Serine 71 was mutated to Alanine (highlighted in yellow) in order to avoid any non specific phosphorylation and binding. Scale bars = 20 µm.

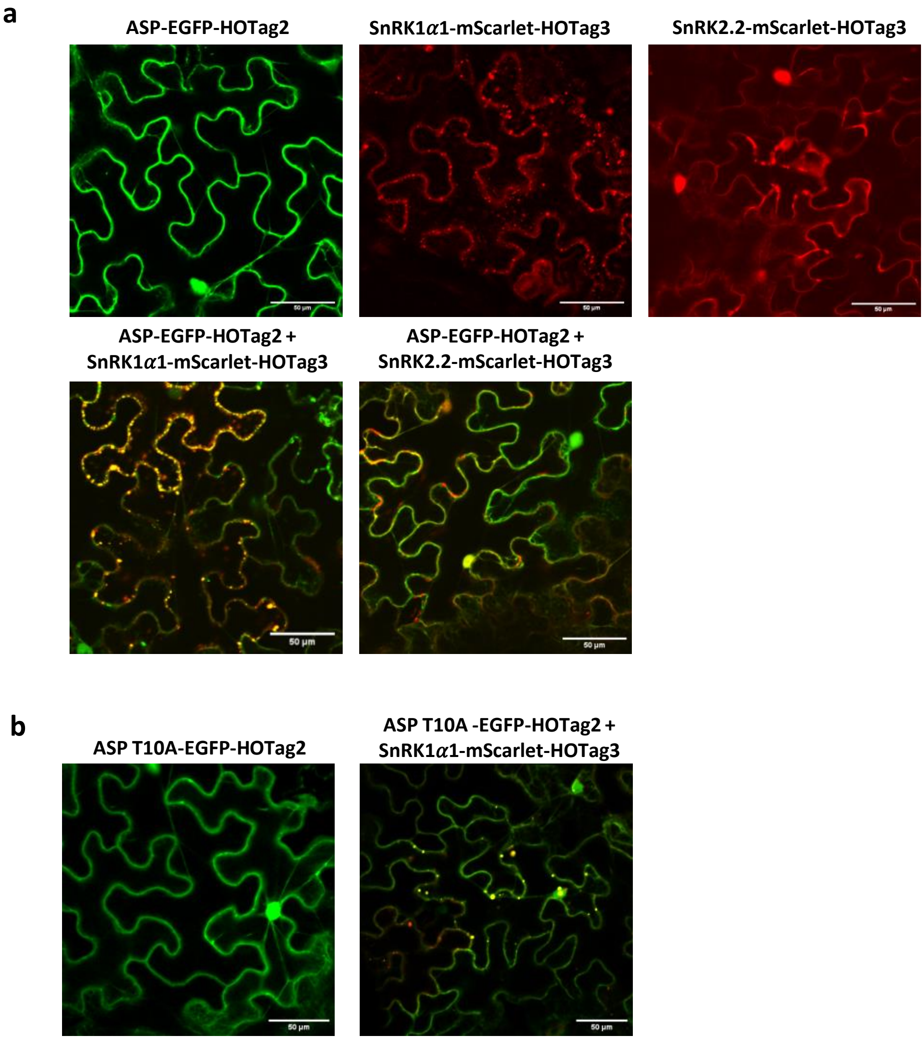


**Supplementary Fig. 7 l ASP peptide interacts with SnRK1α, independently from Thr10. a,** Confocal images of *N. benthamiana* pavement cells transiently expressing either ASP-eGFP-HOTag2 (ASP-HT2), SnRK1α1-mScarlet-HOTag3 (SnRK1α1-HT3) , SnRK2.2-mScarlet-HOTag3 or a combination of ASP-HT2 with one of the SnRK-HTs. **b,** Co-expression of the mutated ASP-T10A-HT2 with the SnRK1α1-HT3 fusions also formed droplets. Scale bar = 50 µm.

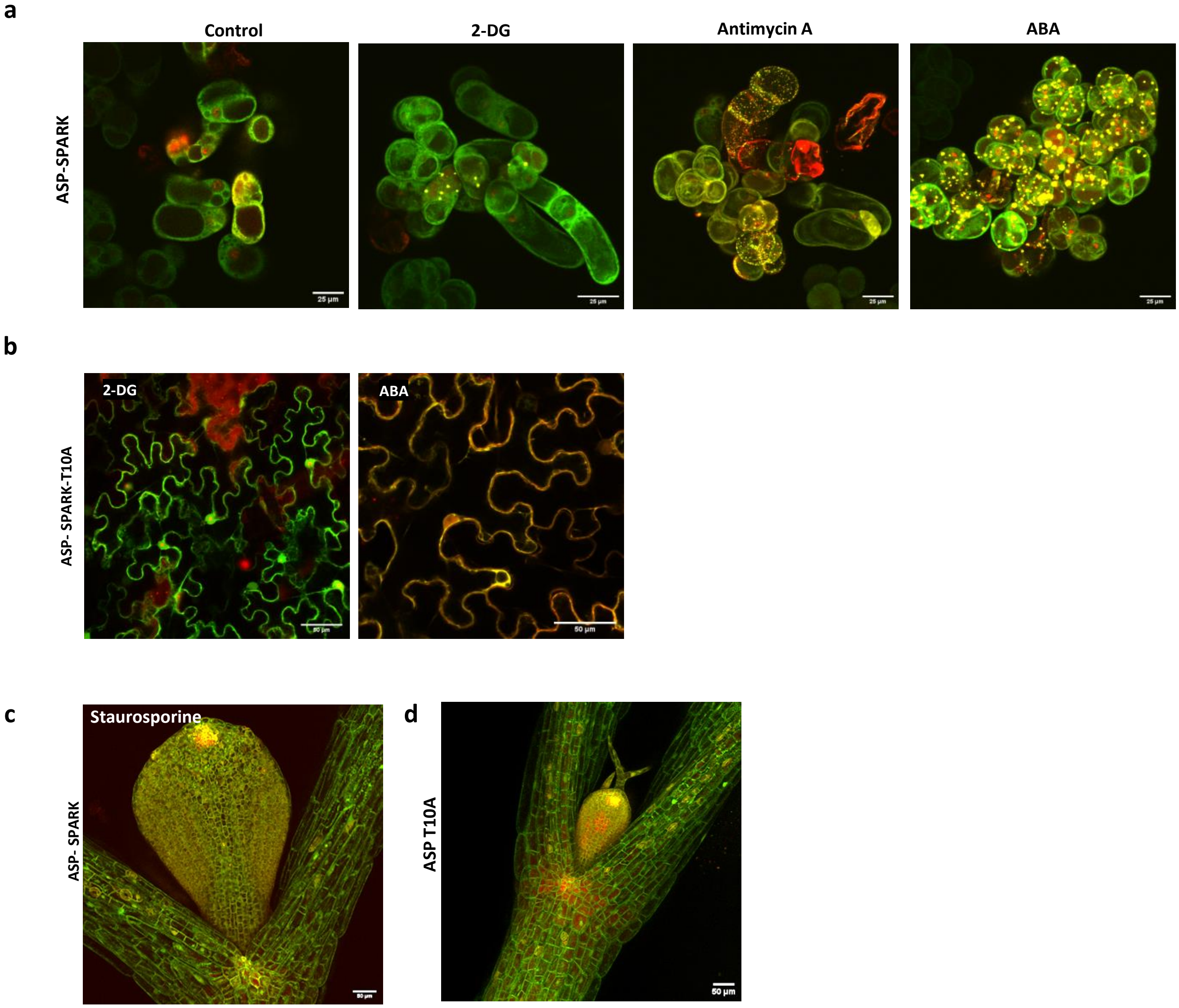


**Supplementary Fig. 8 l a,** PSB-D cells expressing ASP-SPARK before treatment, or 8h after addition of 50mM 2-DG, 20 µM Antimycin A or 10 µM ABA. **b,** ASP T10A-SPARK in *N. benthamiana,* pavement cells treated with 10mM 2-DG or 5 µM ABA for 30 minutes. **c,** ASP-SPARK in *A. thaliana* treated with Staurosporine for 16h (overnight) **d,** ASP T10A SPARK in *A. thaliana.*

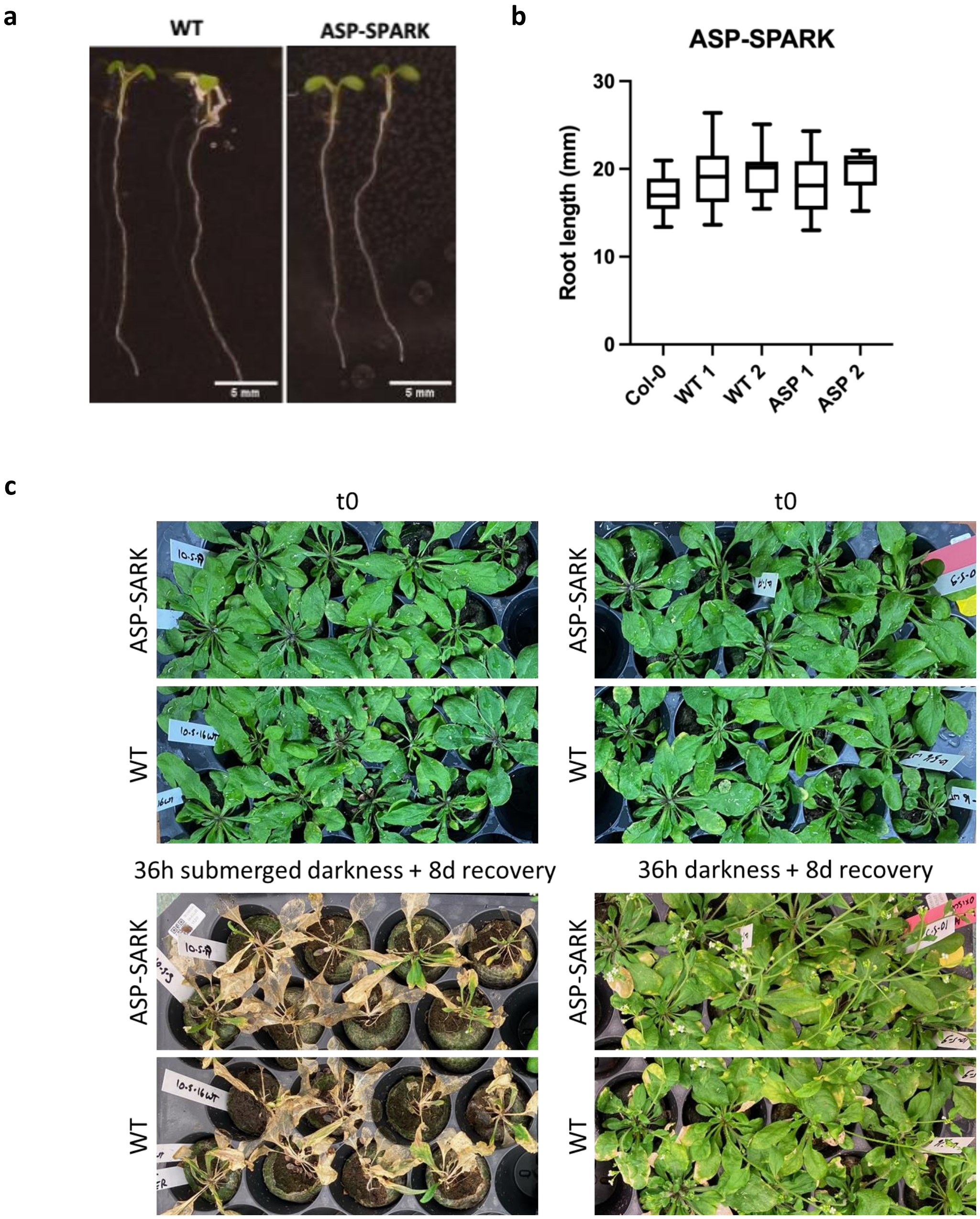


**Supplementary Fig. 9 l a,** ASP-SPARK seedlings compared to WT Col-0 seedlings 7 DAG. **b,** Quantification of root length in from 2 independent ASP-SPARK lines relative to Col-0 and the respective segregating WT siblings at 7 DAG (n=12) **c,** 6 week old ASP-SPARK plants do not show an altered recovery after complete energy deprivation (36h darkness or 36h darkness + submergence) compared to WT plants. $T_0$ pictures were taken just before the onset of the treatment. Survival was scored 8 days after the end of the treatment.
