## Supplementary Table 1 for "Phase separation-based visualization of protein-protein interactions and kinase activities in plants"

**Supplementary Table 1 | Overview of the clones that are used in this study.**

| plasmid | comments |
|---|---|
| pcDNA3-ERK-SPARK 106921 | addgene |
| pcDNA3-PKA-SPARK 106920 | addgene |
| **Gateway vectors** | |
| pDONR221-FkBP12 | |
| pDONR221-FRB | |
| pDONR221-CDKA1 | |
| pDONR221-CKS1 | |
| pDONR221-TPLATE | (Van Damme et al., 2006) [1] |
| pDONR221-LOLITA | (Gadeyne et al., 2014) [2] |
| pDONR221-TASH3 | (Gadeyne et al., 2014) [2] |
| pDONR221-AKS1 | |
| pDONRP2R.P3-EGFP-Hotag6 | |
| pDONRP2R.P3-EGFP-Hotag3 | |
| pDONRP2R.P3-mScarlet-Hotag6 | |
| pDONRP2R.P3-mScarlet-Hotag3 | |
| pcDNA3-FRB-EGFP-HOTag6 | |
| pcDNA3-FKBP12-EGFP-HOTag3 | |
| **Golden Gate entry clones** | |
| PGG-B-ERK-C | ERK's target peptide |
| PGG-B-ERK_T48A-C | |
| PGG-B-SOMA(long)-C | 90 amino acids (MKP1) |
| PGG-B-SOMA(long)_T64A-C | |
| PGG-B-SOMA-C | 15 amino acids (MKP1) |
| PGG-B-SOMA_T64A-C | |
| PGG-B-SNACS-C | 48 amino acids (AKS1) |
| PGG-B-SNACS_S30A-C | |
| PGG-B-WW-C | |
| PGG-B-FHA1-C | |

| | |
|---|---|
| PGG-B-14-3-3-C | |
| PGG-C-EGFP-HOTag3-D | |
| PGG-C-EGFP-HOTag6-D | |
| PGG-D-NOSt-E | |
| PGG-E-35St-F | |
| pGG-F-A-AarI-SacB-AarI-G-G | (Decaestecker et al., 2019) [3] |
| PGGB-FRB+linkerI-C | |
| PGGB-FKBP12-C | |
| PGGB-FHA1-C | |
| PGGB-ASP-C | |
| PGGB-ASP T10A-C | |
| PGGC-LinkerI-EGFP-Linker II-Hotag3-D | |
| PGGD-SPARK6-E | D-LinkerI-EGFP-LinkerII-HOTag6-D |
| PGGD-linkerII-HOTag1-E | |
| PGGD-linkerII-HOTag2-E | |
| PGGD-linkerII-HOTag3-E | |
| PGGD-linkerII-HOTag4-E | |
| PGGD-linkerII-HOTag5-E | |
| PGGD-linkerII-HOTag6-E | |
| PGGD-linkerII-HOTag7-E | |
| PGGA006 (A-pPcubi10-B) | (Lampropoulos et al., 2013) [4] |
| PGGA004 (A-pCAMV35S-B) | (Lampropoulos et al., 2013) [4] |
| PGG-A-pAT1G11910 (pAPA1)-B | (Waadt et al., 2020) [5] |
| PGGC-mCherry-D | (Lampropoulos et al., 2013) [4] |
| PGGC-EGFP-D | (Lampropoulos et al., 2013) [4] |
| PGGE-OCSt-F | |
| PGGE-G7T-F | |
| PGGF KanR | |
| PGGF BaR | |

**Plant destination clones**

| | |
|---|---|
| pKAG KanR | (Decaestecker et al., 2019) [3] |
| pPAG | |
| pK-U1-A-ccdB-G-U9 | Golden Gibson expression vector |
| **Plant expression clones** | |
| UNS2-35S-FHA1-mCherry-H6-OCSt-UNS3 | |
| UNS2-pAPA1-FHA1-mCherry-H6-OCSt-UNS3 | |
| UNS1-35S-ASP-EGFP-H3-OCSt-UNS2 | |
| UNS1-35S-ASP T10A-EGFP-H3-OCSt-UNS2 | |
| UNS1-pAPA1-ASP-EGFP-H3-OCSt-UNS2 | |
| UNS1-pAPA1-ASP T10A-EGFP-H3-OCSt-UNS1 | |
| pK-U9U1 35S:FHA1-mCherry-H6-OCSt-35S:ASP T10A-EGFP-H3-OCSt | Tobacco experiments |
| pK-U9U1 35S:FHA1-mCherry-H6-OCSt-35S:ASP-EGFP-H3-OCSt | |
| pK-U9U1 pAPA1:FHA1-mCherry-H6-G7t-pAPA1:ASP-EGFP-H3-OCSt | PSB-D experiments |

1 Van Damme, D. *et al.* Somatic cytokinesis and pollen maturation in Arabidopsis depend on TPLATE, which has domains similar to coat proteins. *Plant Cell* **18**, 3502-3518, doi:10.1105/tpc.106.040923 (2006).

2 Gadeyne, A. *et al.* The TPLATE adaptor complex drives clathrin-mediated endocytosis in plants. *Cell* **156**, 691-704, doi:10.1016/j.cell.2014.01.039 (2014).

3 Decaestecker, W. *et al.* CRISPR-TSKO: A Technique for Efficient Mutagenesis in Specific Cell Types, Tissues, or Organs in Arabidopsis. *Plant Cell* **31**, 2868-2887, doi:10.1105/tpc.19.00454 (2019).

4 Lampropoulos, A. *et al.* GreenGate---a novel, versatile, and efficient cloning system for plant transgenesis. *PLoS One* **8**, e83043, doi:10.1371/journal.pone.0083043 (2013).

5 Waadt, R. *et al.* Dual-Reporting Transcriptionally Linked Genetically Encoded Fluorescent Indicators Resolve the Spatiotemporal Coordination of Cytosolic Abscisic Acid and Second Messenger Dynamics in Arabidopsis. *Plant Cell* **32**, 2582-2601, doi:10.1105/tpc.19.00892 (2020).
